# Neural signatures of model-based and model-free reinforcement learning across prefrontal cortex and striatum

**DOI:** 10.1101/2025.01.11.632388

**Authors:** Bruno Miranda, James L. Butler, W. M. Nishantha Malalasekera, Timothy E. J. Behrens, Peter Dayan, Steven W. Kennerley

**Author notes:** equal contribution (Bruno Miranda, James L. Butler).

## Abstract

Animals integrate knowledge about how the state of the environment evolves to choose actions that maximise reward. Such goal-directed behaviour - or model-based (MB) reinforcement learning (RL) - can flexibly adapt choice to changes, being thus distinct from simpler habitual - or model-free (MF) RL - strategies. Previous inactivation and neuroimaging work implicates prefrontal cortex (PFC) and the caudate striatal region in MB-RL; however, details are scarce about its implementation at the single-neuron level. Here, we recorded from two PFC regions – the dorsal anterior cingulate cortex (ACC) and dorsolateral PFC (DLPFC), and two striatal regions, caudate and putamen – while two rhesus macaques performed a sequential decision-making (two-step) task in which MB-RL involves knowledge about the statistics of reward and state transitions. All four regions, but particularly the ACC, encoded the rewards received and tracked the probabilistic state transitions that occurred. However, ACC (and to a lesser extent caudate) encoded the key variables of the task - namely the interaction between reward, transition and choice – which underlies MB decision-making. ACC and caudate neurons also encoded MB-derived estimates of choice values. Moreover, caudate value estimates of the choice options flipped when a rare transition occurred, demonstrating value update based on structural knowledge of the task. The striatal regions were unique (relative to PFC) in encoding the current and previous rewards with opposing polarities, reminiscent of dopaminergic neurons, and indicative of a MF prediction error. Our findings provide a deeper understanding of selective and temporally dissociable neural mechanisms underlying goal-directed behaviour.

## Introduction

Animals use at least two major reinforcement learning (RL) systems for behavioural control in sequential decision-making: a goal-directed or model-based (MB) system and a habitual or model-free (MF)^1–4^ system. Both approaches rely on previous experience and converge to the same behaviour given enough practice in a stable environment, but they differ as to how this information is used to infer the values of choices. MB-RL computes estimates prospectively by integrating reward information with knowledge about the state-transition function, which specifies how the state of the world evolves probabilistically given particular choices^5,6^. MF-RL, a less flexible but simpler approach, learns without any model of the environment by bootstrapping sampled experience to train cached predictions of long-run rewards via reward prediction errors (RPEs)^7,8^.

Neural substrates of reward-based learning and decision-making involve complex anatomical connections between the basal ganglia and the prefrontal cortex (PFC)^9,10^. Lesion studies and human neuroimaging suggest an involvement of the primate putamen in MF-RL^11–13^, whereas more anterior regions of the caudate^13–15^, PFC^16–23^ and hippocampus^19,24,25^ have been implicated in MB-RL. However, there is also much work on the online^26–28^ and offline^29–32^ integration of MF and MB quantities, emphasising a closer inter-play between both learning systems. At the single-neuron level, midbrain dopaminergic cells report a RPE that could drive the updating of MF-RL predictions^33,34^. Furthermore, neurons in the striatum have been found to encode cached MF-RL action-values at choice time^35^, but the paradigms used did not elicit differences between MF and MB computations.

One such task that does enable the dissociation of MB and MF computations is Daw et al. (2011)’s ‘two-step’ task^18^. It contains a probabilistic transition between task states to uncouple MF learners (who would assign credit to which state was rewarded regardless of the transition) from MB learners (who would appropriately assign credit based on the reward and transition that occurred). Rodents^19^, monkeys^36^, and humans^18^ all use MB-like behaviour to solve the task. Evidence in rodents suggests dorsal anterior cingulate cortex (ACC) tracks rewards, states, and the probabilistic transition structure, and that ACC is essential in implementing a MB-strategy^37^. Here, we compare primate single neuron activity of 4 different subregions implicated in reward-based learning and choice (ACC, dorsolateral PFC (DLPFC), caudate, and putamen) during performance of the classic two-step task, and demonstrate signatures of MB-RL primarily in ACC, and MF-RL signatures most notably in putamen.

## Methods

### Subjects and neurophysiological procedures

Subjects C/J were two male rhesus monkeys (Macaca mulatta) that were 5-6 years of age and weighed 8–10kg at the time of neural recordings. Surgical procedures were performed using aseptic techniques and under general anaesthesia. Subjects were implanted with a titanium head positioner for restraint, then implanted with two recording chambers that were located based on 3T MRI and stereotactic measurements. The centre of each chamber along the anterior–posterior (AP) coordinate plane was as follows: left hemisphere at AP = 38(C)/37(J) mm, right hemisphere at AP = 27(C)/27.5(J) mm. The chambers were angled along the medial–lateral plane to target different regions. Craniotomies were then performed inside each chamber to allow for neuronal recordings. We used gadolinium-attenuated MRI imaging and electrophysiological mapping of gyri and sulci to confirm chamber placement and electrode trajectories within our recording grid. A custom-built MATLAB® (version R2014b, MathWorks, Massachusetts, USA) algorithm was used to project each recording location (using grid position and depth from dura penetration) onto the MRI images (Figures S1-S2).

Neuronal activity was measured with epoxy-coated (FHC Instruments, Bowdoin, USA) or glass-coated (AlphaOmega Engineering, Nazareth, Israel) tungsten microelectrodes inserted through a guide tube mounted in a custom-designed grid with 1 mm spacing between adjacent grid locations. Electrodes were slowly advanced through the dura each recording session using either custom-built manually controlled micro-drives that lowered electrodes in pairs or triplets, or from motorised microdrives (Flex MT and EPS, Alpha Omega Engineering, Nazareth, Israel) with individual control of electrodes. During a typical recording session, 8–28 electrodes were lowered bilaterally into multiple target regions until well-isolated neurons were found. Neuronal signals were recorded at 40 kHz (OmniPlex System, Plexon Instruments, Dallas, USA). Single-unit isolation was achieved with manual spike sorting (Offline Sorter by Plexon Instruments, Dallas, USA). Neurons were randomly sampled; no attempt was made to select neurons based on responsiveness or specific cortical layer. After each recording session, the microelectrodes were retracted and the microdrive assemblies were removed.

We recorded neuronal data from four target regions: ACC, DLPFC, caudate and putamen (Figures S1-S2). In subject C, we recorded simultaneously from the ACC (dorsal bank of the ACC sulcus, primarily area 9/32) and the DLPFC (dorsal bank of the principal sulcus, area 46d) in both the left and right hemispheres, and from the dorsal caudate and the dorsal putamen in the right hemisphere. In Subject J, we recorded from the ACC (dorsal bank of the ACC sulcus, primarily area 9/32), and the DLPFC (dorsal bank of the principal sulcus, area 46d) in the left hemisphere, and from the dorsal caudate and the dorsal putamen from the right hemisphere. In total we recorded single-unit activity from 661 neurons (C: 508 and J: 153) in 57 recording sessions (C: 30 and J: 27) across all four investigated regions: ACC: 240 neurons, DLPFC: 187 neurons, Caudate: 115 neurons, Putamen: 119 neurons. In some sessions, neurons were recorded from all four regions simultaneously, whereas in other sessions, only two or three regions were sampled.

### Task

Full task details are reported elsewhere^36^. We monitored eye position and pupil dilation during the task using an infra-red system (ISCAN ETL-200). In brief, subjects initiated a trial by fixating a central red cue, and once extinguished, subjects were free to view the stimuli and indicate their decision by moving a manual joystick in the direction of the chosen stimulus (Figure 1). Two decisions were required (first-stage, second-stage states) on each trial to obtain reward. The first-stage state was indicated by a grey background and the choice was between two options indicated by pictures. Each first-stage choice could lead to either a common (70% transition probability) or rare (30% transition probability) second-stage state, represented by different background colours (brown and violet). This state-transition structure was fixed throughout the experiment. In the second-stage state, another two-option choice between pictures was required, which could lead to three different outcome levels: high (big reward and no delay), medium (small reward and small delay) or low (no reward and big delay). To promote learning and updating of stimulus values, each second-stage stimulus had an independent reward structure: the outcome level (high, medium, low) remained the same for 5-9 trials, and then, either stayed the same level (with one-third probability) or changed randomly to one of the other two possible outcome levels. Rather than fixed reward amounts, small Gaussian drifts of reward (mean/standard deviation of 0/200ms for high reward and 0/100ms for medium reward) were also added to promote constant valuation of the reward amounts. Fifteen percent of the trials were forced (i.e., without allowing a choice as only one option was presented), which could be at either the first or second-stage. The trial type sequence was randomly generated at the start of the session and was followed even after error trials. Trials with either no choice, no eye fixation, break of eye fixation, early joystick response, or the joystick not centred before choice resulted in time-outs for the subjects, and were excluded from the data analysis (C: M = 5%; J: M = 8%). In both decision stages, the choice options were randomized to one of three possible locations, and subjects indicated their choice by moving a joystick towards the stimulus (C: left, right and down; J: left, right and up). The reward (C: diluted cranberry juice; J: diluted apple juice) was provided by a spout positioned in front of the subject’s mouth and delivered at a constant flowrate using a peristaltic pump (Ismatec IPC). We used Monkeylogic software (http://www.monkeylogic.net/) to control the presentation of stimuli and task contingencies and acquire joystick and eye data. All visual stimuli used were the same across sessions for both subjects. All experimental procedures were approved by the UCL Local Ethical Procedures Committee and the UK Home Office (PPL Number 70/8842) and carried out in accordance with the UK Animals (Scientific Procedures) Act.

**Figure 1.**
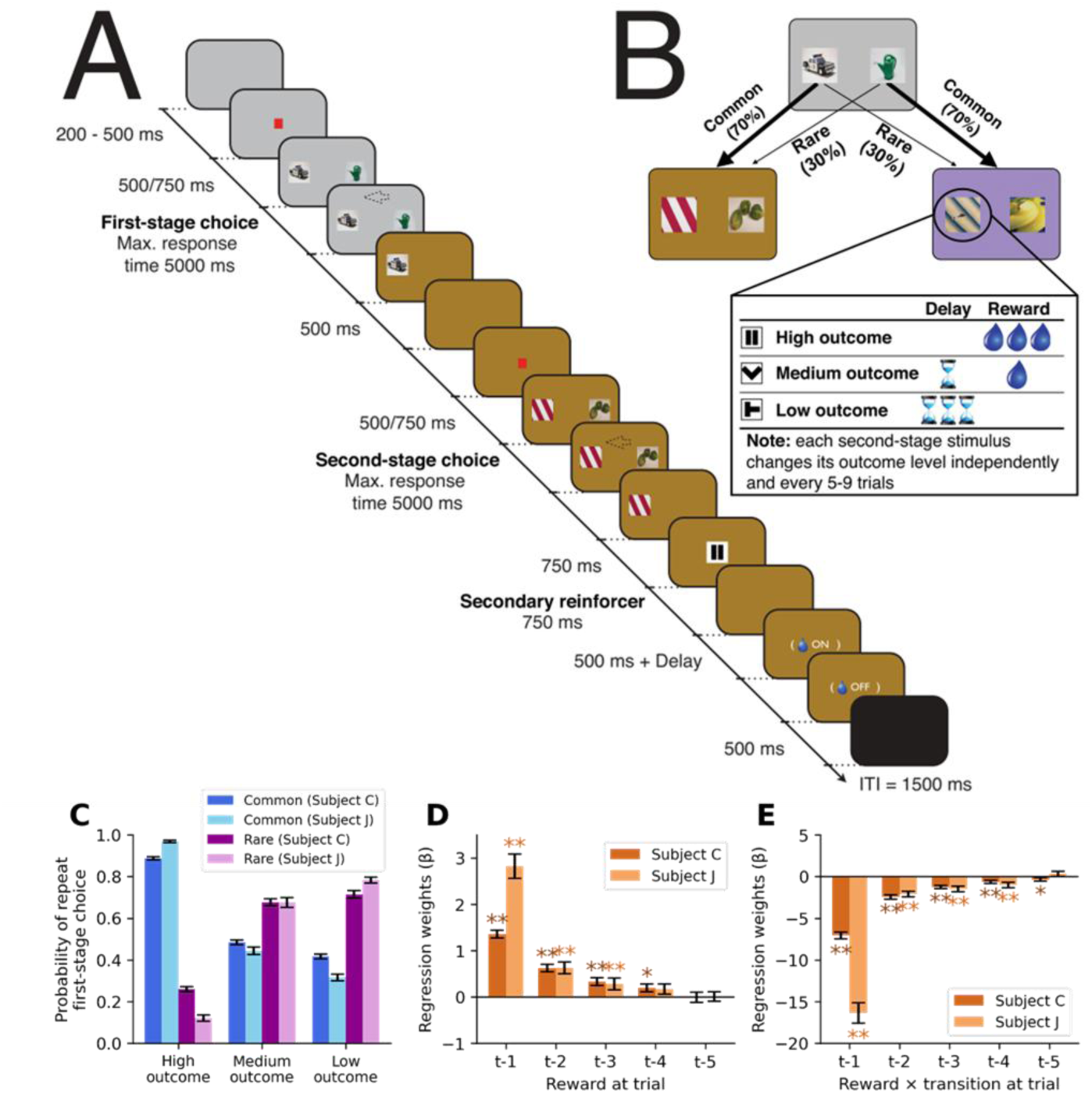
Two-stage decision task performance. **A**, Timeline of events. Eye fixation was required while a red fixation cue was shown, otherwise subjects could saccade freely and indicate their decision (arrow as an example) by moving a manual joystick in the direction of the chosen stimulus. Once the second-stage choice had been made, the nature of the outcome was revealed by a secondary reinforcer cue (here, the pause symbol represents high outcome). Once the latter cue was off the screen, there was a fixed 500 ms delay and the possibility of a further delay (for both medium and low outcomes) before juice was provided (for both high and medium outcomes). **B**, The state-transition structure (kept fixed throughout the experiment). Each second stage stimuli had an independent reward structure: the outcome level (defined by the magnitude of the reward and the delay to its delivery) remained the same for a minimum number of trials (a uniformly distributed pseudorandom integer between 5 and 9) and then, either stayed in the same level (with one-third probability) or changed randomly to one of the other two possible outcome levels. **C**, Likelihood of first-stage choice repetition, averaged across sessions, as a function of reward and transition on the previous trial. **D-E**, Logistic regression results on first-stage choice with the contributions of the reward main effect (**D**) and reward × transition (**E**) from the five previous trials. **, *, p < 0.05, 0.01, respectively. In **C-E**, error bars depict standard error of the mean.

### Neural analysis

All data analysis was conducted using Python 3.7.6 (Python Software Foundation).

#### Raster generation

Each cell’s spike raster was smoothed with a gaussian kernel (σ = 50ms) and epoched relative to the time at which either an epoch started (choice 1, choice 2, feedback) or an action was taken (initial fixation, choice 1 made, choice 2 made). Rasters were then downsampled to 100Hz resolution by averaging every 10 points and standardised using the mean and standard deviation of each epoch across all trials.

#### Multiple linear regression models

Multiple linear regression was used to index encoding of different task parameters. Three different general linear models (GLMs) were constructed. The first design matrix (GLM1) included task parameters without considering any algorithmically derived MF or MB computations (Table 1). The second design matrix (GLM2) included state value estimates of choice 1 and choice 2’s value derived from the best-fitting computational models which generate MB and MF Q-value estimates on each trial^36^ (Table 2).

**Table 1.** The contents of GLM1. Far right column indicates which figure panels each regressor was used for.

| Regressor | Description | Figure |
| --- | --- | --- |
| Constant | To account for the mean effect |  |
| Linear effects | A ramping regressor to account for any monotonic autocorrelations across the session |  |
| (Linear effects) ^ 2 | To account for any polynomial autocorrelations across the session |  |
| (Linear effects) ^ 3 |  |  |
| (Linear effects) ^ 4 |  |  |
| Sin 2 | Sin curves of varying frequency (approximately 2, 4, 6, 9 cycles per session) to account for any non-monotonic autocorrelations across the session |  |
| Sin 4 |  |  |
| Sin 6 |  |  |
| Sin 9 |  |  |
| Previous reward (t-3) | Value of the secondary reinforcer shown three trials ago |  |
| Previous reward (t-2) | Value of the secondary reinforcer shown two trials ago | 2F-G |
| Previous reward (t-1) | Value of the secondary reinforcer shown on previous trial | 2A-C<br>2F-G |
| Previous transition | Transition that occurred on the previous trial |  |
| Previous choice 1 | The first choice from the previous trial |  |
| Previous reward * previous transition | Pairwise interactions between the events from the previous trial |  |
| Previous reward * previous choice 1 |  |  |
| Previous transition * previous choice 1 |  |  |
| Previous reward * previous transition * previous choice 1 | Three-way interaction of the events from the previous trial (equivalent to previous second stage state * previous reward, as choice 1 * transition = second stage state) | 6B |
| Reward | Value of the secondary reinforcer (current trial) | 2D-E<br>3E<br>5E-F |
| Transition | Transition that occurred (current trial) | 3A-E |
| Choice 1 | The cue the subjects chose (current trial) | 6A |
| Previous reward * transition | Pairwise interactions between the current trial's events with reward from the previous trial | 3C-D |
| Previous reward * choice 1 |  |  |
| Transition * choice 1 |  |  |
| Previous reward * Transition * choice 1 | Three-way interaction of the current trial's events with the previous trial's reward |  |

**Table 2.**
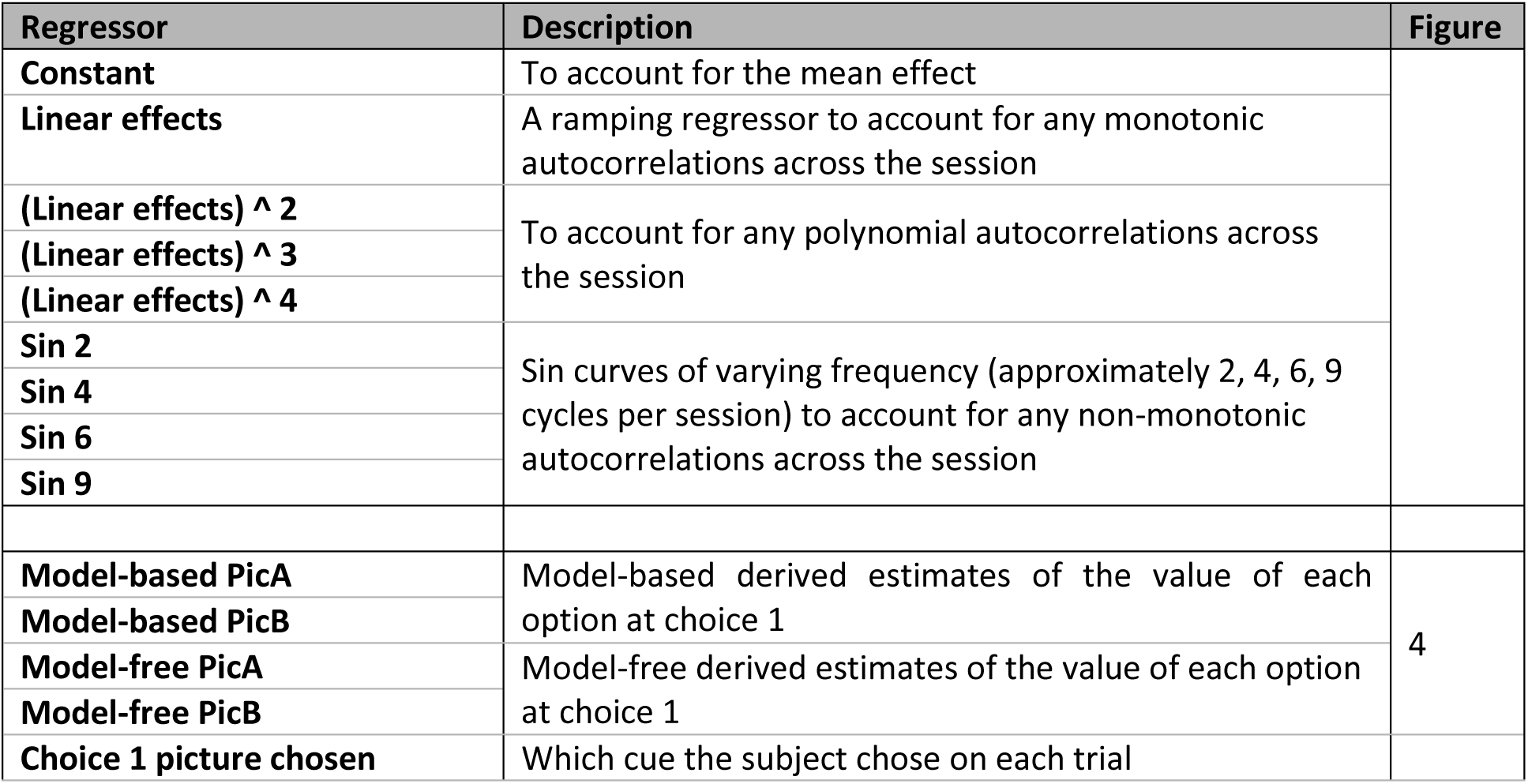
The contents of GLM2. Far right column indicates which figure panels each regressor was used for.

The third design matrix (GLM3) combined MB and MF Q-value estimates based on the animal’s behaviour, which was shown to better predict animal’s choices than any tested MB or MF model in isolation^36^ (‘hybrid’ model) (Table 3).

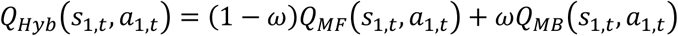

**Table 3.** The contents of GLM3. Far right column indicates which figure panels each regressor was used for.

| Regressor | Description | Figure |
| --- | --- | --- |
| Constant | To account for the mean effect |  |
| Linear effects | A ramping regressor to account for any monotonic autocorrelations across the session |  |
| (Linear effects) ^ 2 | To account for any polynomial autocorrelations across the session |  |
| (Linear effects) ^ 3 |  |  |
| (Linear effects) ^ 4 |  |  |
| Sin 2 | Sin curves of varying frequency (approximately 2, 4, 6, 9 cycles per session) to account for any non-monotonic autocorrelations across the session |  |
| Sin 4 |  |  |
| Sin 6 |  |  |
| Sin 9 |  |  |
| Hybrid-based chosen option | Hybrid-based derived estimates of the value of each option at choice 1 | 5 |
| Hybrid-based unchosen option |  |  |

Equation 1. The Hybrid model assumes that first-stage choices are computed as a weighted sum of the state-action values from MF and MB learning systems. a, action, s, stage (choice 1/2), t, trial, w, fitted hyperparameter.

The GLMs also included several control regressors to account for any neural drift that occurred over time. This included linear and quadratic time functions, and sin curves of various frequency (see Tables 1-3). Examination of the false positive rates using session shuffled data^38–40^ demonstrated that these adequately controlled for neural drift and autocorrelations (Figure S3).

### Population level encoding

To assess the presence of coding for a particular explanatory variable in multiple linear regression we used the coefficient of partial determination (𝑝𝑎𝑟𝑡𝑖𝑎𝑙 𝑅^2^, CPD):

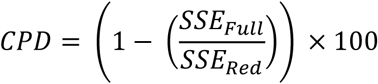

Where SSE is the sum of squared error using either the full design matrix (SSE_Full_) or the design matrix with the relevant regressor omitted (SSE_Red_). This represents the percentage of additional variance that the full model explains compared to the reduced model. This measure was then averaged across all cells from a region and the mean and standard error used as summary statistics.

To determine significance of our CPD measure we computed the null distribution by shuffling the dependent variable 500 times and repeating the analysis. Several control regressors were also included to capture autocorrelation in the data (Table 1-3). Rare cases where NaNs were generated due to a lack of variance in certain time points from low firing rate neurons were discarded. Periods of encoding that crossed the 95^th^ percentile of this distribution were then considered as putatively significant. A second null distribution was then computed of the lengths of periods of significance that occurred in the null distribution. Significant events in the true data had to remain above threshold for longer than the Bonferroni corrected (by 4 as there were 4 areas) 95^th^ percentile of the null distribution of lengths to be counted as significant. This resulted in an acceptable false positive rate of approximately 5% (Figure S3).

For comparing correlations between weights for different features (i.e., between transition and reward coding, Figure 3E), the null distribution of correlations observed in circularly shifted data was compared to the correlation seen in the actual data. This accounts for any correlations between features that existed in the task by preserving the structure of the design matrices.

To compare differences in CPD between regions, an independent sample’s t-test was initially used to find periods where CPD’s differed significantly. A permutation test was then conducted by shuffling neurons between the two regions and repeating the t-test over time for 500 permutations. Any observed runs of significance in the real data had to be longer than the 95^th^ percentile of runs of significance in the null distribution.

### Single neuron encoding

An equivalent cluster-based permutation test was used to quantify the number of individual neurons that significantly encoded a particular parameter. The neurons firing rate in the relevant epoch was shuffled across trials and the null distribution of the CPD computed for 500 permutations. The neuron’s CPD then had to be higher for 95% of the null distribution, for longer than 95% of runs of spurious significance in the null distribution, to be considered significant. This resulted in an acceptable false positive rate of approximately 5% (Figure S3).

To test whether the proportion of significant neurons was larger than expected by chance, a binomial test against 5% was used. To test whether there were significant regional differences in the proportion of significant neurons observed, a chi-square test was used.

### Pattern of encoding

Similarities in the pattern of neural encoding between different parameters was quantified using Pearson’s r. Significance was determined by computing the null distribution of correlations by shuffling coefficients. Any observed runs of significance had to be longer than 95% of any spurious runs of significance that occurred in this null distribution.

### Decoding analysis

A support vector machine (SVM) was used to decode the first-stage choice (choice 1). First, a pseudopopulation was generated by collapsing across all neurons recorded from a region. The session with the fewest number of trials for each sample was determined, and all other sessions randomly subsampled to this length to create a dataset with an equal number of samples for each neuron and each condition. The data were then split using stratified five-fold cross validation; whereby 80% of trials were used to train an SVM (using sklearn.svm.LinearSVC from the Python Scikit-learn package), which was then tested on the remaining 20% of trials. This process was repeated for the remaining 4 folds of the data and decoder accuracies averaged over the 5 folds. As this process involved leaving out a random selection of trials each time, it was repeated 20 times and the average and standard error of these 20 permutations logged as the indicator of the decoder’s performance.

To determine whether the decoder was performing significantly better than chance, the above procedure was repeated 500 times while shuffling the trial labels on each iteration to ascertain the null distribution of the decoder’s performance. The observed data were then compared to this distribution to calculate the reported p-values, and significance counted as an accuracy > 95% of the null distribution. To compare whether one condition was significantly better than another, a null distribution of differences was created by shuffling condition labels and calculating the difference in decoder performance between the randomly split conditions 500 times. The observed difference in accuracy between conditions was then compared to this null distribution of differences.

## Results

Two male rhesus macaques (Macaca mulatta) completed a variation of the classic two-step task^18^. After an initial eye fixation period, subjects first chose between two first-stage options (A, B, ‘choice 1’), each leading to a different second-stage state (indicated by background colour) (Figure 1A, B). Crucially, the transition between these two stages was probabilistic, with each option transitioning to its preferred second-stage state in 70% (‘common’) of trials; and transitioning to the alternative second-stage state in 30% (‘rare’) of trials (Figure 1B). This state-transition allows the dissociation of MF-RL strategies (that estimate state values using bootstrapped sampling of experienced states) from MB-RL strategies (which use the transition structure to assign rewards prospectively to states based on the probability of that state occurring). Both second-stage states then required another choice (‘choice 2’) between two options (C and D, or E and F), leading to one of three possible rewarding outcomes (high, medium, low outcome) indicated by a secondary reinforcer cue (‘feedback’). In our version of the task, the outcome values of each second-stage state randomly and independently changed value every 5-9 trials, thus requiring subjects to continue sampling the different second-stage states to determine where the highest rewards were located across trials (Figure 1B). This reward schedule resulted in an advantage of a MB-policy over an MF-policy, the former of which consistently gained more reward per trial than the latter in simulations [MB gained 1.2979 ± 0.0071 units of reward per trial (n=57 recording sessions, the 3 reward levels were counted as 0, 1, or 2 AU); MF 1.2416 ± 0.0069 reward per trial; difference between them: p<0.0001, paired t-test; Figure S4]. Both strategies were also better than a random policy (0.9953 ± 0.007 reward per trial, vs MF/MB: p<0.001).

In a prior comprehensive behavioural analysis^41^, we demonstrated that both subjects estimated the value of each choice 1 option using a combination of MF and MB-RL algorithms, but with MB dominating (Figure 1C-E). MF-RL does not exploit information about task structure, so it predicts no difference in the probability of repeating choice 1 dependent on common/rare transition, whereas a key signature of MB-RL is just such a difference. We found that both subjects were significantly more likely to repeat choice 1 when a high reward was obtained through a common (rather than rare) transition, with the opposite pattern evident following low rewards (Figure 1C), indicating a strong MB influence on behaviour. Having established the influence of MF- and MB-RL strategies on behaviour^36^, we next examined whether signatures of MF- and MB-RL were evident in neuronal activity in four key brain regions implicated in learning and choice: we recorded single-neuron activity from two regions in PFC – dorsal anterior cingulate cortex (ACC; n=240) and dorsolateral PFC (DLPFC; n=187), and two regions in the striatum – caudate (n=115) and putamen (n=119) (see Figures S1-S2 for recording locations).

As parameters were correlated over trials (e.g., the same choice was made repeatedly when it led to a high reward), there was a risk that intrinsic temporal autocorrelations in firing rate across trials would confound the analysis^38–40^. We therefore incorporated several nuisance regressors to account for temporal autocorrelations in neuronal activity that may have been present (see Methods, Tables 1-3). To check this was an adequate control, we analysed neural firing rates with respect to a different session’s trial data (i.e., the dependent and independent variables for the regression were from different sessions)^38–40^. No significant encoding of any of our parameters of interest were observed at either the population or single neuron level in such null controls (Figure S3). Therefore, temporal autocorrelations in neuronal activity were adequately controlled, and we next turned to examining neural encoding of task parameters.

### ACC encoded reward across task events; striatal regions encoded a MF RPE signal

We first examined the extent of reward encoding in each region by examining the response of neurons to the secondary reinforcer cue. At the population level, all regions strongly encoded the value of the secondary reinforcer cue at feedback; furthermore, this encoding persisted throughout a vast period of the following trial – being significantly stronger for ACC during this particular time period (Figure 2A; GLM1). The same was true at the single neuron level, with large populations of neurons within each region encoding the reward during feedback and long into the following trial (Figure 2B). ACC’s encoding of reward was stronger than all other regions at both the population level (p<0.05 cluster-based permutation test; Figure 2A) and single neuron level (p<0.05, chi-square test; Figure 2B). Encoding of the value of the secondary reinforcer peaked earlier in striatal neurons (507.7 ± 19.16-ms) compared to prefrontal neurons (588.5 ± 13.9 ms; p=0.005 independent t-test; Figure 2C). Therefore, all four regions, but especially ACC, encoded the previous trial’s reward throughout the next trial.

**Figure 2.**
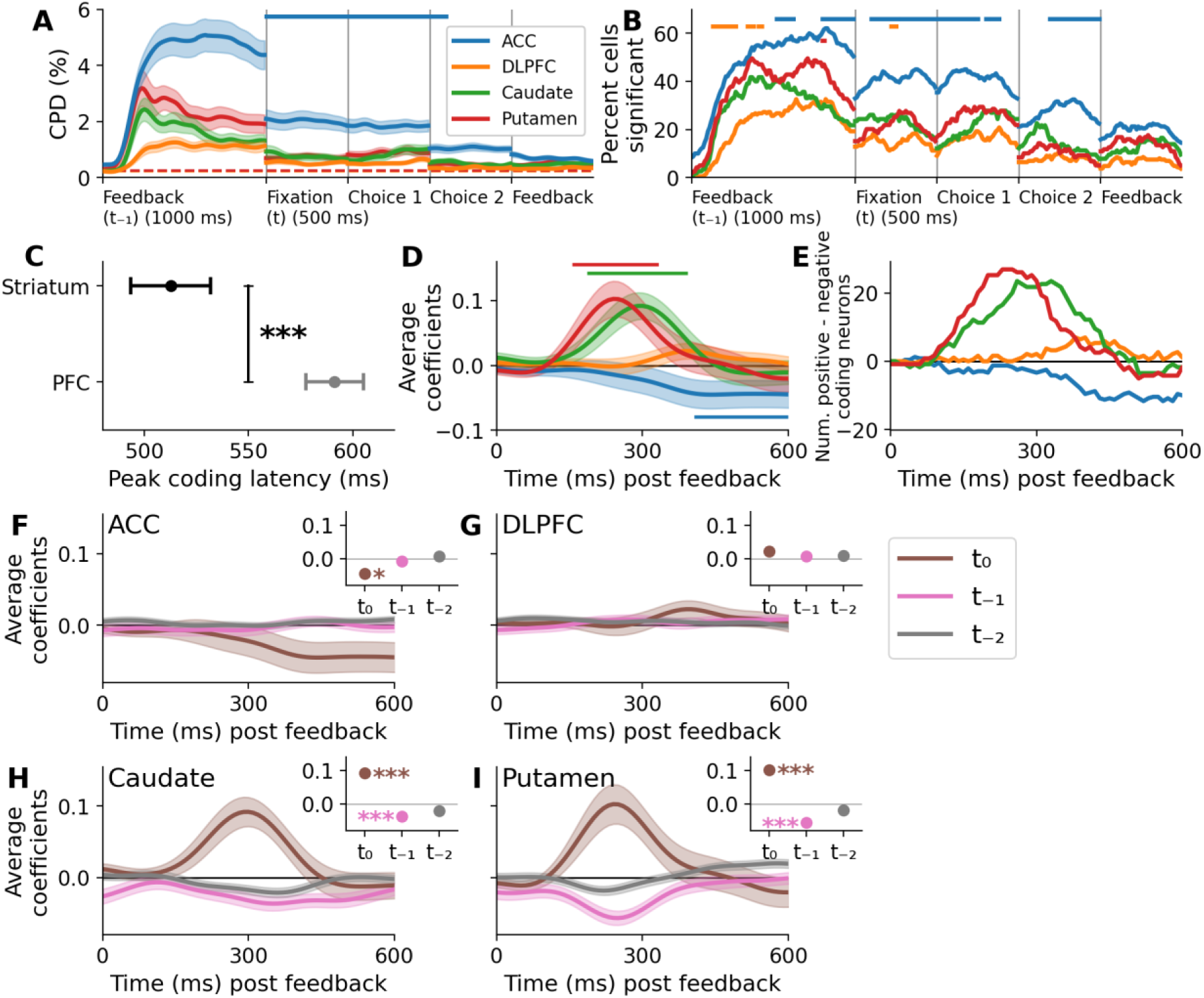
ACC is the strongest reward coding region. **A**, Average coefficient of partial determination (CPD) across neurons in each region for the encoding of the reward received on the previous trial. Feedback is shown both for the previous trial (left) and the current trial (right). Solid horizontal lines represent periods where that area’s CPD differed significantly from all other areas CPDs (p<0.05, cluster-based permutation test). Dashed horizontal line indicates the confidence interval for each region derived from the null distribution. All epochs are 0-500ms with the exception of 0-1000ms for the first feedback epoch. **B**, Percent of neurons within each region that significantly encoded reward in the feedback epoch and epochs of the subsequent trial (p<0.05 cluster-based permutation test). Solid horizontal line represent period where that area’s CPD differed significantly from all other areas CPDs (p<0.05, chi-squared test). **C**, The time (relative to feedback onset) at which the peak encoding of the secondary reinforcer occurred, across significant encoding neurons. ***,0.001, independent t-test. **D**, Average coefficients for encoding the value of the secondary reinforcer. Solid horizontal lines represent periods where the coefficients in a particular region differed from 0 (p<0.05, 1-sample t-test). **E**, Same as in **D** but for the percentage of significant neurons. Dashed lines indicate the net percentage of significant neurons (positive – negative percentages). **F**, Average coefficients across ACC neurons for the reward received on the current trial and the previous two trials. Inset, the average activity for each of the three conditions at the time point of peak reward encoding. **, ***, p<0.01,0.001 1-sample t-test against 0. **G-I**, Same as **F** but for DLPFC, Caudate, and Putamen neurons, respectively. In all cases, error bars depict standard error of the mean across neurons.

The direction of this reward coding differed between regions from PFC and striatum. The latter encoded the predicted reward (i.e., from secondary reinforcer) positively (i.e., firing rate positively correlated with reward; p<0.05 from 190- and 160-ms for caudate and putamen respectively, 1 sample t-test of coefficients against 0; Figure 2D, E); whereas ACC encoded reward with a negative polarity and slower response time (p<0.05 at 410-ms onwards, Figure 2D, E). Additionally, both caudate and putamen encoded the reward from the previous trial negatively during the feedback period of the current trial (Figure 2F, G). Such quantitative features in both striatal areas (i.e., positive coding of current reward and negative encoding of past rewards) is typical of a dopaminergic RPE signal^33,34^. ^3^

**Figure 3.**
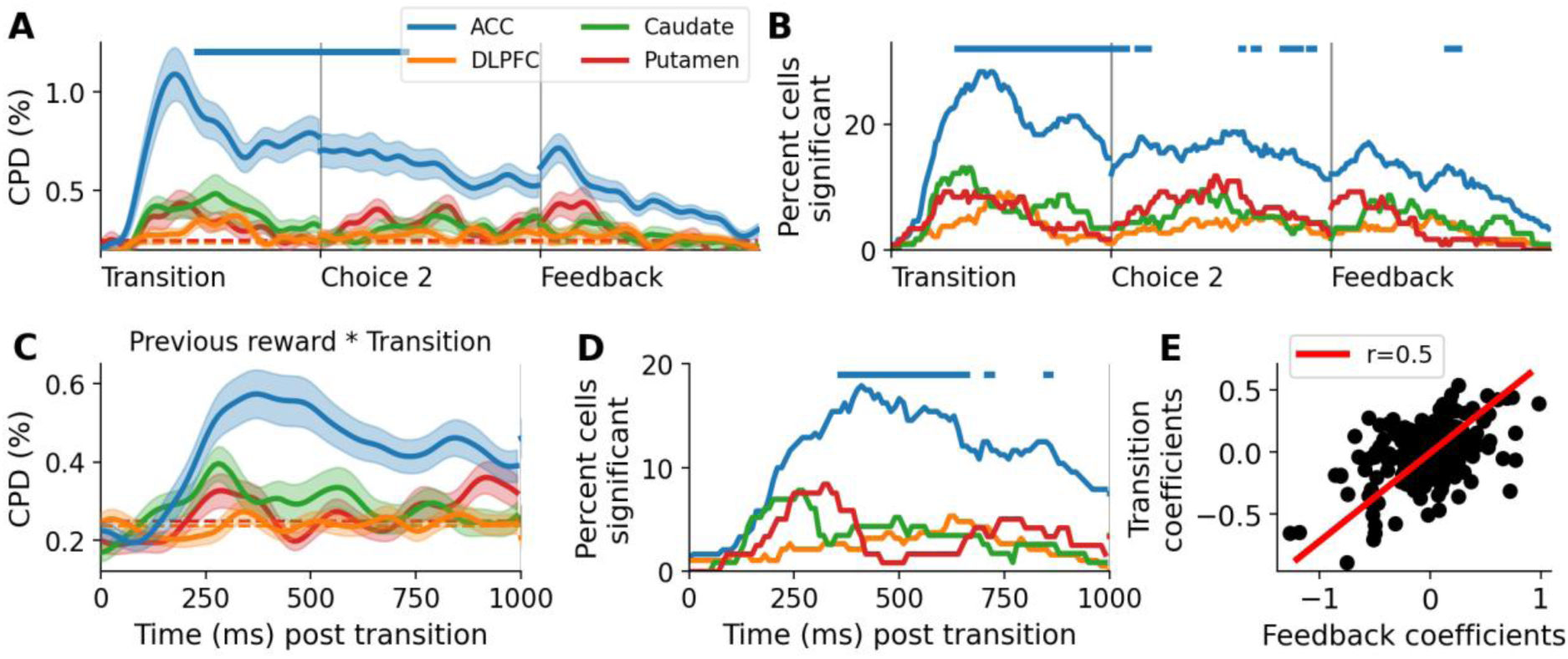
ACC encodes transition information until feedback. **A**, Average coefficient of partial determination (CPD) across neurons in each region for the encoding of the transition that occurred. Solid horizontal lines represent periods where that area’s CPD differed significantly from all other areas CPDs (p<0.05 cluster-based permutation test). Dashed horizontal line indicates the confidence interval for each region derived from the null distribution. All epochs are 0-1000ms. Error bars depict standard error of the mean. **B**, Percent of neurons within each region that significantly encoded transition (p<0.05 cluster-based permutation test). **C**, **D**, Same as **A, B**, but for encoding of the interaction of previous reward and transition. **E**, ACC coefficients for transition and feedback were correlated (coefficients taken from 300 ms post onset of each epoch). In **A**, **B**, and **D**, solid horizontal line indicates periods where ACC was significantly greater than all three other regions (**A**: permutation test, **B** and **D**: chi-squared test, p<0.05/3).

### ACC encoded MB state-transition information

Tracking the state-transition structure of the task is imperative for solving the task as a MB-learner. All four regions encoded whether the current trial’s first-stage choice transitioned to the common or rare second-stage state (which could be inferred by a change in background colour immediately after choice indicating which second stage state they had just entered, Figure 1A). This signal was significantly stronger in ACC – where state-transition information was maintained from transition until the feedback epoch (CPD > 0.3%, p<0.002, cluster-based permutation test; Figure 3A; GLM1). A similar pattern was also observed at the individual neuron level (Figure 3B).

As reward expectations would change on rare transitions if using a MB strategy, we next examined whether the encoding of previous trial reward was modulated by current trial’s transition. While all four regions exhibited some selectivity for the interaction between the previous trial’s reward and the current trial’s transition, a larger proportion of ACC neurons encoded this parameter compared to all other regions (p<0.05, chi-squared test, Figure 3C-D). Thus, ACC reward expectancy signals were modulated by the state-transition structure of the task.

All regions, but particularly ACC, encoded a common transition (at the time of transition) similar to a high reward (at the time of feedback), as there was a positive correlation between the coefficients for reward and transition (the transition parameter was signed such that common and rare transitions were equivalent to high and low rewards, respectively) (ACC r=0.4963, DLPFC r=0.3273, caudate r=0.4712, putamen, r=0.5052; all p<0.002 except DLPFC where p=0.006, circular permutation test; Figure 3E, S5). As the reward expectation will be higher on common compared to rare trials, this demonstrates that the brain encodes being diverted to an area with a lower reward expectation equivalent to actually receiving a low reward (and vice versa). This signal also reflects awareness of the (MB-like) state-transition structure.

### ACC and Caudate encoded MB-value estimates

Due to this encoding of the state-transition and its interaction with other variables, we next tested whether neurons encoded MB- or MF-derived chosen values of the choice 1 options (using RL computational modelling to estimate these values in a subject- and trial-specific manner^36^). At the single neuron level, ACC was unique in having a large population of neurons that encoded both the MB and MF value of the choice 1 options (Figure 4A-B; GLM2). The encoding of MB emerged 280 ms before the cues were displayed, indicating the subjects were anticipating the upcoming choice, whereas MF-derived chosen value coding only emerged after the cues were presented (170 ms post cue presentation, p<0.05 binomial test). Caudate also had a population of neurons encoding the MB-derived value of choice 1, which emerged 430-ms after choice onset (Figure 4A). At the population level all 4 regions encoded the MB value estimate of choice 1, with ACC and caudate encoding it more than 500-ms before the options were displayed (Figure 4C). All 4 regions, also encoded MF-RL estimates of choice 1’s value, again with ACC encoding emerging more than 500-ms before choice onset (Figure 4D).

**Figure 4.**
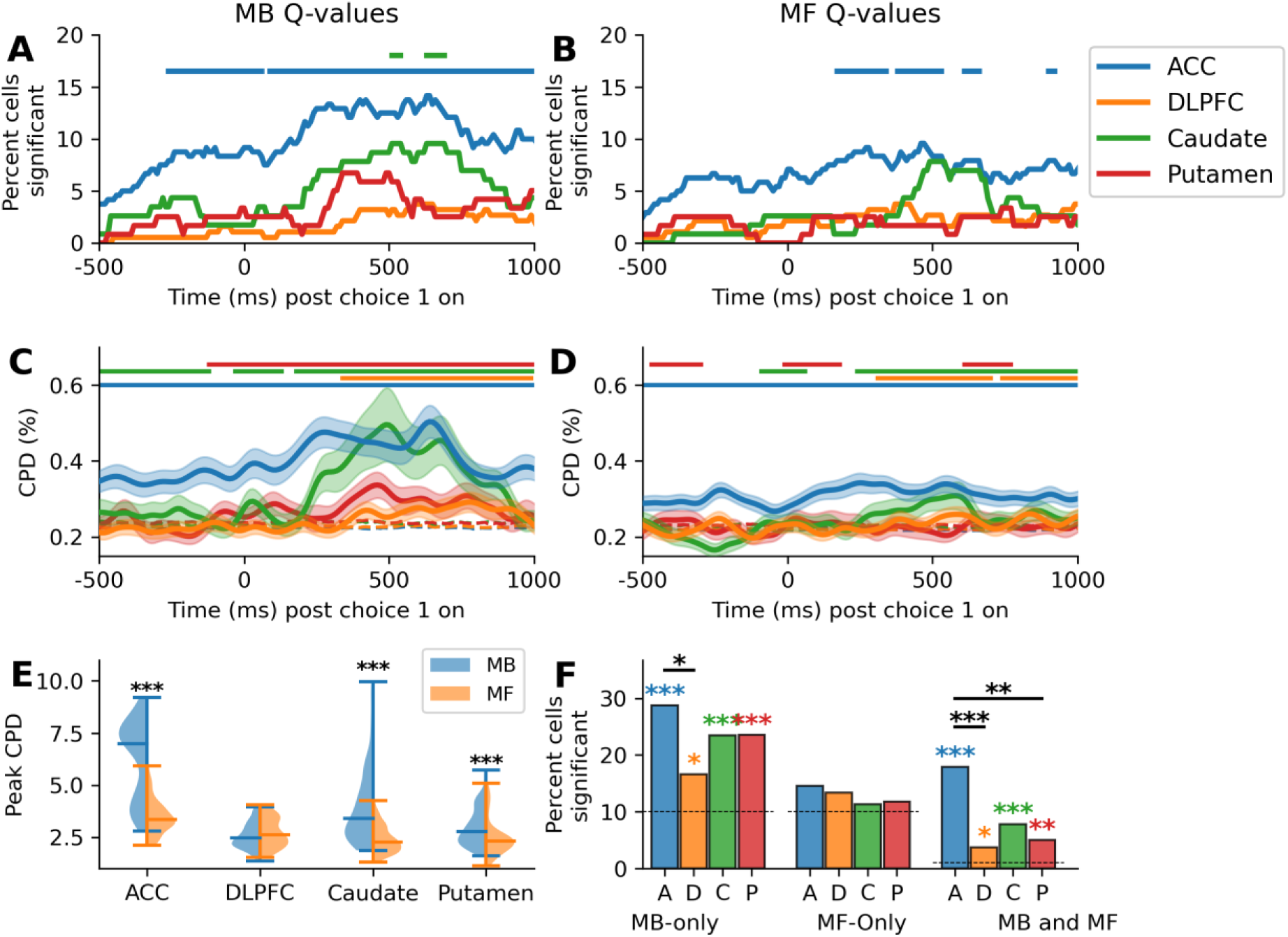
Value estimate encoding was predominantly model-based. **A**, Percentage of cells in each region that encoded the model-based (MB) derived estimates of the value of each of the choice 1 options. Solid horizontal lines indicate periods where the percentage of neurons was significant (p<0.05, binomial test). **B**, Same as in **A**, but for model-free (MF) derived estimates of each option’s value. **C**, **D**, Same as in **A**, **B** but for the average CPD across neurons in a region. Dashed lines represent the 95% confidence interval determined by permutation testing. Solid lines indicate periods where the strength of encoding was significant (p<0.05, cluster-based permutation test). Error bars depict standard error of the mean. **E**, Distribution of the peak CPD values (i.e., the highest CPD value observed over the epoch shown in A-D for either PicA or PicB cues) for each neuron during the epoch shown in **C** (MB, blue) and **D** (MF, orange) values. Horizontal lines indicate median and extrema. *, ***, p<0.05,0.001 paired t-test. **F**, The percentage of neurons in each region that significantly encoded a MB-estimate of the chosen option’s value (left), a MF estimate (middle), or both (right) assessed using cluster-length permutation testing. Coloured asterisks indicate that population is significantly greater than 10% (blue and orange) or 1% (green), binomial test. Asterisks between bars indicate a difference in size between the two populations (*, ***, p<0.05, 0.001, chi-square test).

To further compare this encoding, we examined the peak strength of encoding of each neuron for either of the two choice 1 options (i.e., MB and MF value estimates for PicA/PicB; Table 2) at any point from 500-ms before to 1000-ms after choice 1 was presented. In all regions except DLPFC, peak CPD values across neurons were higher for MB compared to MF estimates (ACC: t=19.8812, p<0.0001 paired t-test; DLPFC: t=-2.1474, p=0.1319; caudate: t=14.659, p<0.0001; putamen: t=6.1911, p<0.0001; Figure 4E). ACC’s peak MB and MF encoding was significantly higher than in all other regions (p<0.001 independent t-test).

Examination of the strongest signal observed, ACC’s encoding of MB Q-values, showed a dynamic pattern with different neurons encoding the signal at different parts of the epoch (Figure S6). When aggregating the number of significant coders throughout the epoch, and examining the specificity of MB versus MF coding, we found that all regions had a significant population of neurons that encoded MB-, but not MF-, derived value (30, 19, 23 and 24% of neurons in ACC, DLPFC, caudate and putamen respectively; all p<0.0014 binomial test against 10% (as the strongest response to either of the two options was used); Figure 4F). All regions had a population of neurons that encoded both MB- and MF-derived option value (18, 4, 8, 5% of neurons in ACC, DLPFC, caudate and putamen respectively; all p<0.012 binomial test against 1; Figure 4F), but ACC’s population was significantly higher than all other regions (p<0.05, chi-square test, Figure 4F). This dominance of MB encoding in caudate but not putamen at the population level may explain similar dissociations in these two striatal regions observed in human neuroimaging studies^13^.

### Caudate value estimates remapped following a rare transition

The observed prominence of transition-related selectivity, in addition to strong encoding of MB-derived value estimates, led us to next examine how transition type altered the value estimation of the chosen first-stage option. To explore this, we used a combination of MB- and MF-value estimations (or the hybrid estimate) derived from the animal’s behaviour, which better explained their choices during the task than either of the two models in isolation^36^ (GLM3). In common trials, caudate was unique in encoding the value of the chosen option more positively than the unchosen option’s value (p<0.025 280- to 320-ms post transition revealed, paired t-test; Figure 5). After a rare transition, caudate was again unique in encoding the chosen and unchosen option values differently, but this time it was the unchosen option that was encoded positively and the chosen option negatively (p<0.025 170- to 370-ms, paired t-test; Figure 5). Thus, caudate’s encoding of an option’s value also reflected the availability of the option.

**Figure 5.**
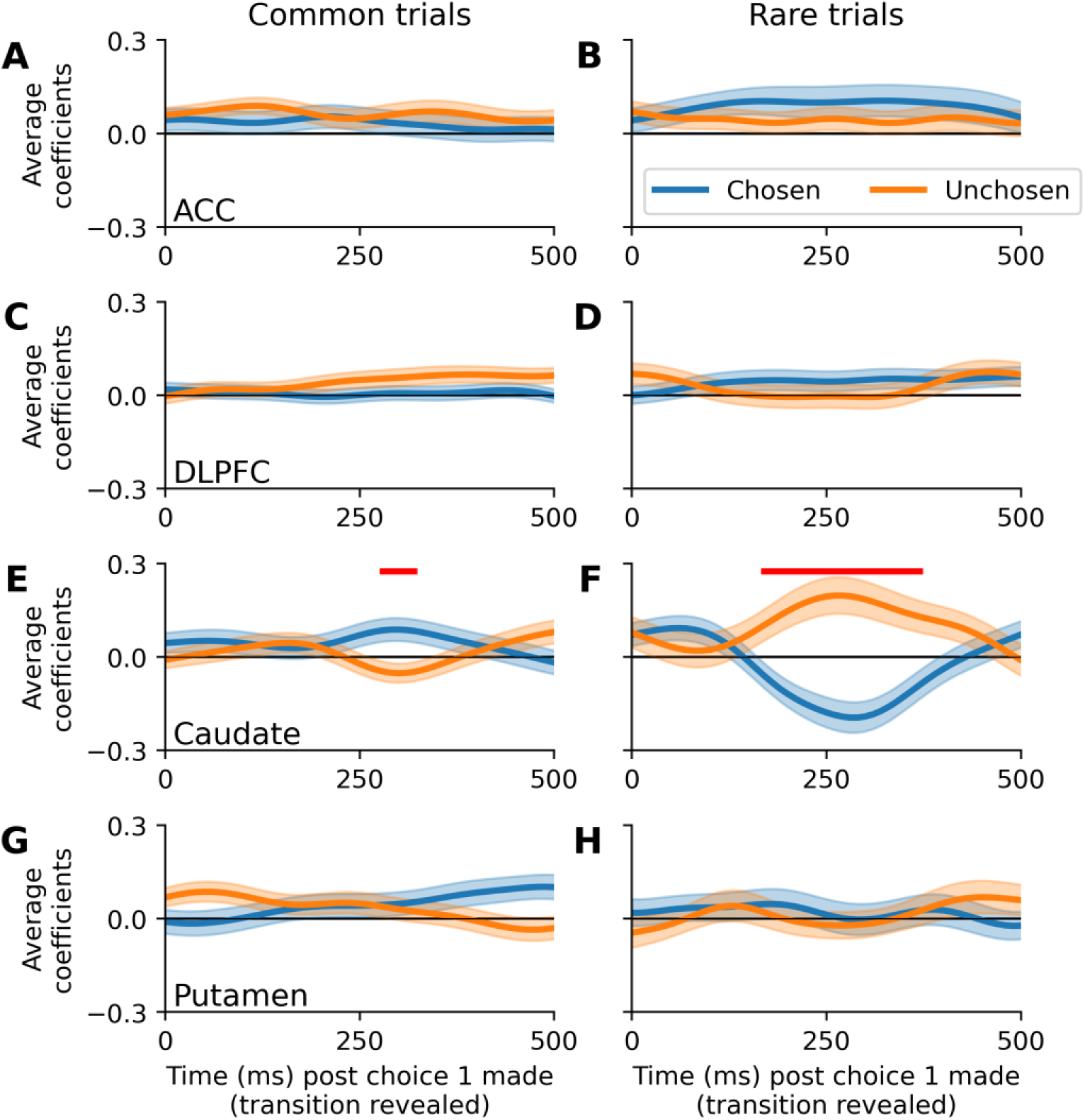
Direction of the encoding of chosen and unchosen choice 1 options, depending on the transition that occurred. **A**, Average coefficients across neurons in the ACC with respect to the value of chosen (blue) and unchosen (orange) options of choice 1 in common trials. Value estimates were calculated using a hybrid of MB and MF estimates derived from each monkey’s behaviour. Red horizontal line indicates portions where the coefficients in the two conditions differed significantly from one another (p<0.05, paired t-test). Error bars depict standard error of the mean. **B**, Same as in **A**, but for trials where a rare transition occurred. **C-H**, Same as in A-B but for DLPFC, Caudate, and Putamen, respectively.

### Reward-modulated encoding of first-stage choice

We next examined the encoding of the choice 1 option (i.e., the stimulus identity of what they chose). All regions encoded it to an equal extent around the time of choice (difference between regions all p>0.05, cluster-based permutation test; Figure 6A; GLM1). Strikingly, all regions also encoded the interaction of choice 1, transition, and reward from the previous trial during the choice 1 epoch (Figure 6B); with ACC encoding it stronger than DLPFC and Putamen (ACC vs DLPFC p=0.006 between -180 to 190-ms, vs Putamen p=<0.002 between -150 to 450-ms; cluster-based permutation test; Figure 6B). Importantly, none of the intermediate pairwise interactions were encoded to a similar extent (Figure S7). Therefore, both ACC and caudate performed a very specific MB computation of integrating the reward, transition, and choice from the previous trial into a signal which could inform which option to choose on the current trial.

**Figure 6.**
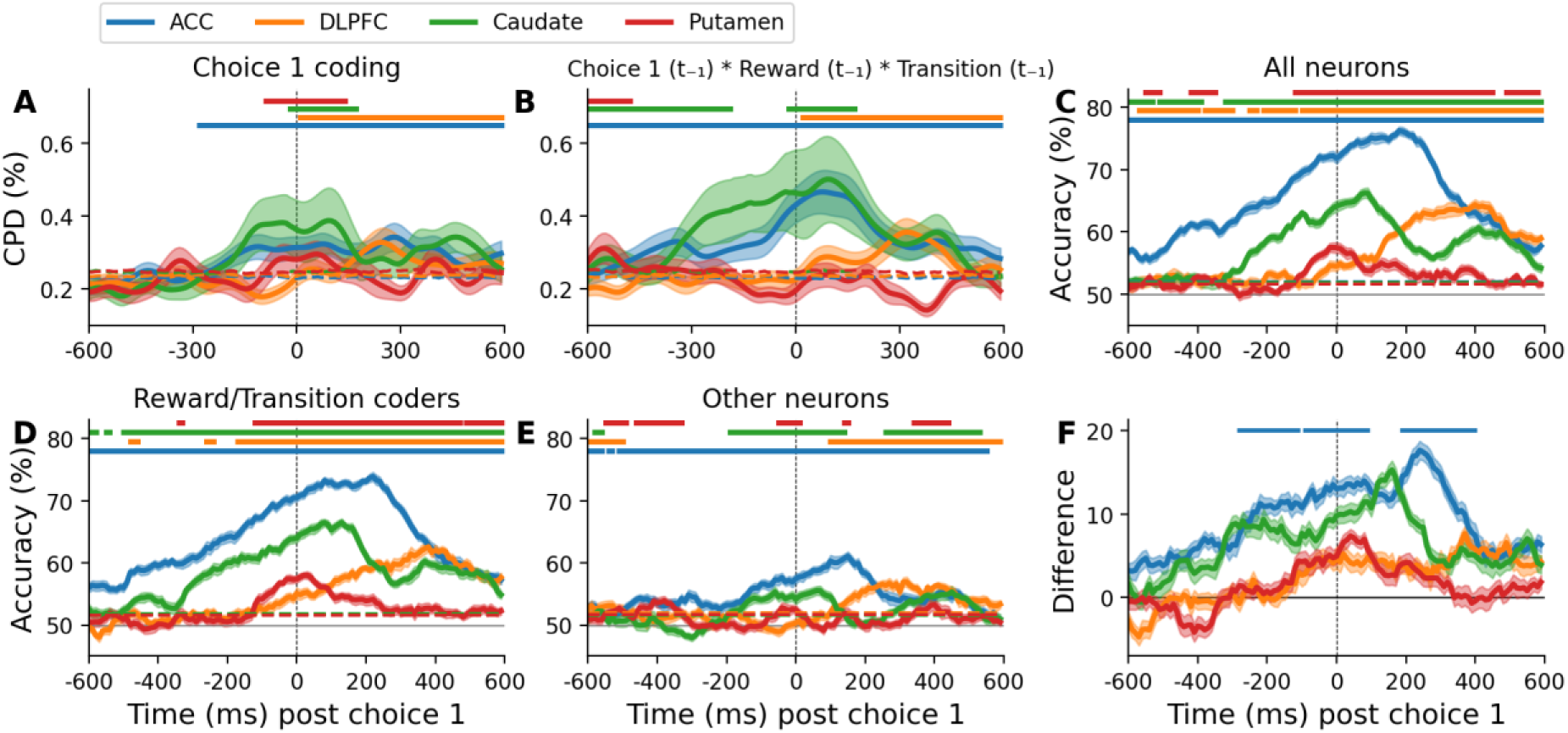
Choice 1 was encoded by neurons sensitive to reward and transition. **A**, Average coefficient of partial determination (CPD) across neurons in each region for the encoding of choice 1. Dashed horizontal line indicates the confidence interval for each region derived from the null distribution. **B**, Same as **A** but for encoding of the interaction of reward, transition, and choice 1, all from the previous trial. **C**, A support vector machine was used to decode from each neural population which cue the monkeys would choose at choice 1 on each trial. Dashed horizontal line indicates the 95^th^ confidence interval (permutation test) and solid horizontal lines indicate periods of significant decoding (p<0.05, cluster-based permutation test). Dashed vertical line indicates the time at which the subjects made their choice (0 ms). **D, E**, Same as in **C** but neurons were median split into two groups depending on the strength with which they encoded reward at feedback and transition at transition. **F**, Difference in decoder strength between **A** and **B**. Solid horizontal line indicates periods of significant difference assessed using permutation test. In all cases, error bars depict standard error of the mean.

Furthermore, we also found that choice 1 stimulus identity was encoded by the same neurons that encoded reward and transition. Choice 1 could be decoded from all four neural populations using a support vector machine, with it being most strongly represented in ACC and caudate (Figure 6C). We then performed a median split of neurons within each region depending on the strength with which they encoded reward at feedback, and transition at transition (taking the average of the two coefficients). In ACC only, the neurons that were more sensitive to reward and transition encoded choice 1 more strongly than those neurons that were less sensitive to these attributes (Figure 6D-F). Therefore, it was the same subpopulation of neurons within ACC that tracked the different task parameters relevant for guiding MB choice.

### Explore/Exploit strategy modulates encoding of first-stage choice

In our task, the outcome level (high, medium, low) of each second-stage stimulus remained the same for 5-9 trials before potentially changing. This design naturally created periods where subjects could ‘exploit’ the same Choice 1 to maximize reward for several trials; and other periods where they had to ‘explore’ different second-stage stimuli to optimize reward (as contingencies shifted). In classical MB-RL, the transition between reward states can be learned by keeping counts of observed transitions from a current state-action pair to a subsequent state, yielding a maximum-likelihood estimate of the environment’s dynamics^42^. In fact, knowledge about the reward contingency schedule could support decision-making in both exploitation – by enabling efficient choice when rewards are stable; and exploration – by guiding alternative behaviour most likely to yield improved outcomes (this is different from MF learning, where exploration is more random since the agent lacks explicit state-transition knowledge).

We thus repeated our decoding analysis of choice 1 stimulus identity, but this time limited trials to those where they had not received a high reward for the previous two trials (‘explore’ trials), and those where the previous two rewards had been the highest level (‘exploit’ trials). All regions encoded choice 1 for some duration of the choice epoch for both explore (p<0.002 in all cases, permutation test; Figure 7A) and exploit (p<0.002 in all cases; Figure 7B) conditions, but decoding accuracy was strongest in ACC. Choice 1 was less strongly decoded – particularly in ACC – in the former condition compared to the latter (p<0.002 in all cases, permutation test on differences observed; Figure 7C); and, also during exploitation, the ACC encoded choice 1 before the choice was even presented to the subject (Figure S8). This pre-choice ACC encoding in exploit trials may reflect the need to allocate cognitive (or attentive) resources to features – i.e., choice 1 stimulus identity – that are most certain predictors of important outcomes. As a control, we also decoded the direction of the Choice 1 (where choice was indicated via joystick movement), which was randomised each trial and therefore orthogonal to the stimulus that was chosen. Again, all four regions encoded its direction in both explore (p<0.002 in all cases; Figure 7D) and exploit (p<0.002 in all cases; Figure 7E). However, there were minimal differences in the strength of the representation between explore and exploit conditions (ACC, p=0.088, cluster-based permutation test; DLPFC p=0.016; caudate p=0.32; putamen p=1; Figure 7F). Therefore, exploit behaviour specifically upregulated relevant task parameters that were worth remembering across trials.

**Figure 7.**
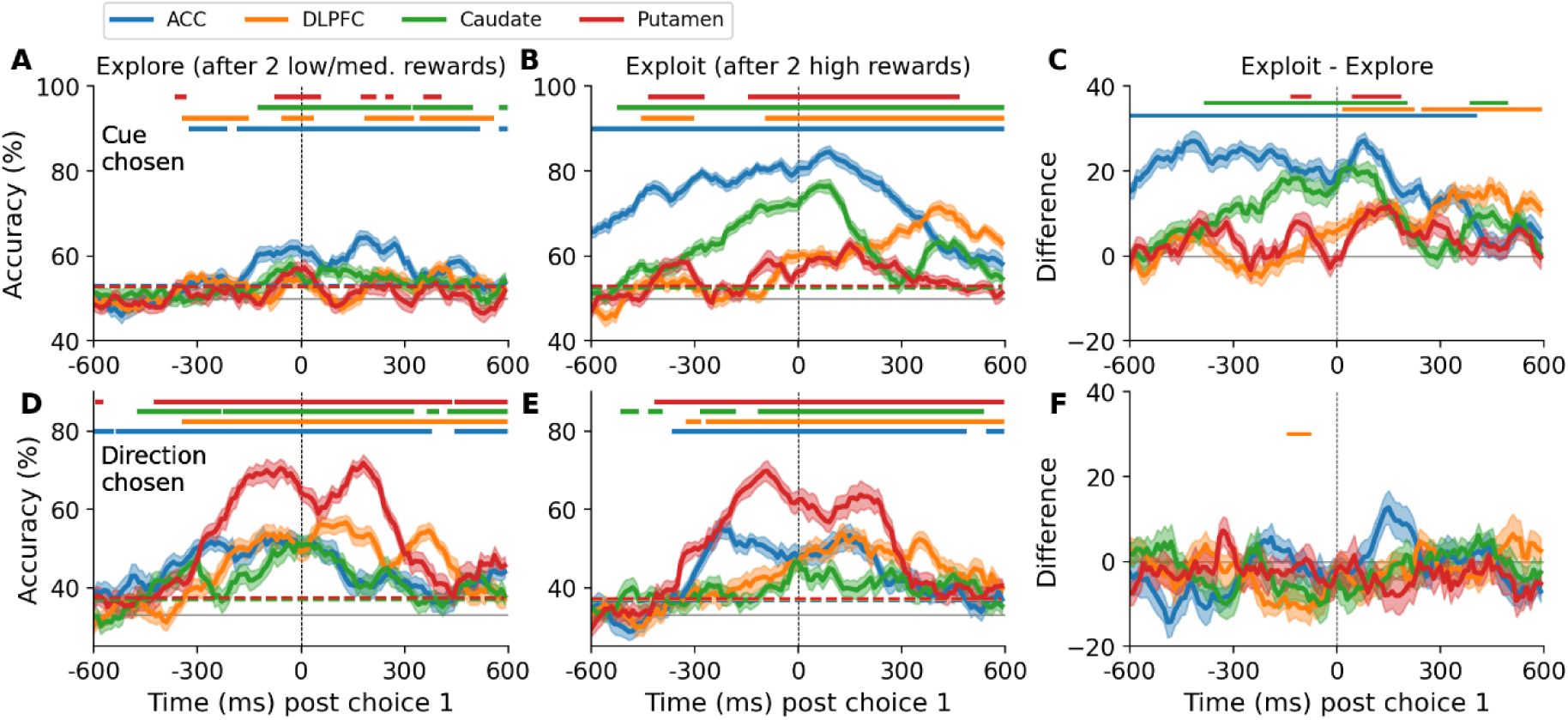
Encoding of choice 1 was modulated by reward stability (explore/exploit strategy) **A**, Trials were split into those following two consecutive high rewards (‘exploit’) and those following two consecutive low/medium rewards (‘explore’). A support vector machine was used to decode from each region which cue the monkeys would choose at choice 1 on explore trials. Shaded error bars represent standard error of the mean over the different permutations of the data. Horizontal dashed lines indicates the 95% confidence interval (permutation test) and solid horizontal lines indicate periods of significant coding (p<0.05, permutation test). **B**, Same as in **A** but only for exploit trials. **C**, Difference in decoder strength between **A** and **B**. Solid horizontal line indicates periods of significant difference assessed using permutation test. **D-F**, Same as in **A-C** but for decoding the direction of the choice 1, which was randomised across trials and therefore orthogonal to the cue that was chosen. Note that a choice was indicated via manual joystick movement.

## Discussion

Despite the extensive work in both MF-RL (or habitual) and MB-RL (or goal-directed) behaviour^3^, few studies have shown simultaneous single-neuron signatures of both learning strategies^37^, let alone across a number of relevant cortical and subcortical regions, and in the primate brain. Using a decision task which formally distinguishes MF and MB values, and by adopting an analytical approach that considered both observable variables and computational measures, we showed that key RL elements underlying choice behaviour were encoded in neurons in the primate PFC and striatum at different time points. The observed widespread and simultaneous representations of MF and MB related computations are consistent with the view that these controllers operate in parallel^2,3,18^, but their associated signals are more richly intertwined than had originally been expected. Additionally, we found convincing evidence supporting a prominent role of ACC, and to a lesser extent, the caudate, in MB-RL. The ability of single neurons to encode reward-related parameters has been described throughout the brain^43–47^. Here we found reward coding at feedback (of MF and MB relevance) to be significantly stronger in ACC. However, at feedback a RPE (a hallmark of MF-RL) can also be computed to support learning. The midbrain dopamine neurons have been strongly implicated in encoding such RPE compared to other areas with similar (but not identical) feedback-related activity^33,34^. Signals in other brain regions, such as PFC, have been found to report differences between received and expected outcome^48–51^, but in most of these cases either the definition of RPE did not conform to the normative RL theory or the neurons did not exhibit the quantitative properties and negative aspect of the error. In our study, both caudate and putamen neurons clearly responded at feedback with the parametric features of a dopamine-like RPE^33,34^. It is important to underline that, despite being seen as a signal that drives MF-RL, both MF and MB approaches coincide at feedback and use this RPE to update their valuations^36^. The data presented here confirms that basal ganglia structures (both dopaminergic and striatal neurons) are unique (relative to PFC) in computing RPE-like signals.

The knowledge of the state-transition structure differentiates MB from MF-RL and hence we investigated neurons that significantly discriminated a common from a rare transition. Our findings are in line with the neural changes seen in the rat ACC when an animal’s belief is modified after an environmental change^52^; and the observation that inhibition of mouse ACC impairs the use of the transition, but not rewards^37^. Additionally, the selectivity of ACC neurons for detecting state-transition fits well with the neuroimaging signal observed in ACC on a saccadic planning task when an internal model required update^53^, endorsing its role in dynamically updating behavioural policies. As all regions adjusted their value estimates as a function of the transition type occurred, our simultaneous recordings extend previous findings by supporting that the ubiquitous use of state-transition information across fronto-striatal circuits is likely propagated from (or via) ACC to other (reward-focused) regions.

It is important to highlight that MB-like behaviour can arise from sophisticated MF-strategies, such as tracking state-transition probabilities and the reward function^37,54,55^. Further work should be undertaken to carefully evaluate whether the MB-like signals seen here are truly derived from a MB-RL mechanism, or instead reflect predictions about hidden states that can be used as inputs for learning by MF systems^55^.

It is well established that ACC neurons can multiplex task variables during decision-making tasks^51,56,57^. The interaction between reward and transition coding is particularly critical for MB-RL valuation and choice in the two-step task. Here we demonstrate that ACC was the primary region to simultaneously encode key variables of the task structure for MB choice, namely the interaction between reward, transition and choice. This was also borne out in the computationally derived MB estimates of the first-stage choice, where ACC activity (and to a lesser extend caudate) strongly encoded this information. Another clear distinction between ACC and the other regions when coding either reward or transition information was the timescale of responses. ACC neurons have some of the slowest intrinsic timescales (i.e. the rate of their autocorrelation decay) across the brain^58,59^, which likely underlies the persistent encoding of these task variables. Maintaining these representations over long time periods likely potentiates the ability to contextualise outcomes by state, as well as the monitoring of action-outcome relationships relevant for behavioural adjustment^60–62^. In fact, the encoding of the next trial’s chosen cue and its expected value was evident in ACC long before the choice epoch even started. Thus, our data provide novel correlates of prospective planning in single-neuron activity^63^ and contributes to our understanding of the mechanisms underlying MB-RL.

Along with ACC, caudate appears to have an important role in MB behaviour. Choice 1 decoding was most strongly represented by ACC and caudate, and specifically by neurons that tracked both reward and transition information. Furthermore, the distinctive caudate signal of updating (flipping) the value estimates of the currently experienced option on rare trials implies this signal is not a general temporal-difference RPE, but instead more reflective of a state-PE^17^, and further supports the role of caudate in MB valuation. Our recordings were predominantly in anterior parts of caudate, which is where others have described flexible value coding^64^, representations of sequences^65^, and preference for early phases of learning^66^ – features more often embraced by MB-RL. Anatomically, it is also the part of the caudate with the highest afferent projections from ACC – the area we observed to support MB computations^67^.

On the other hand, MF-based estimates were neither as striking nor as specific to striatal regions as expected and observed in previous studies^18^. The monkeys were extensively trained on the task before recordings commenced, which may have caused a shift towards both MB behaviour and MB value representation within the striatum. Alternatively, this training may have allowed more sophisticated representations to occur, such as using latent states to expand the task space^54^. Despite this, ACC was the only region that significantly encoded both MB and MF values. Having access to both MF and MB values at the time of choice, implicates ACC in the arbitration process between the two learning strategies^68^ that has been considered crucial to guide optimal behaviour. It is also in line with alternative theoretical proposals for ACC that propose a specific role in monitoring the error likelihoods of all possible expected outcomes^69^. In fact, the use of a RL framework could indeed unify other roles attributed to ACC, in particular conflict resolution^70^ and cognitive control^71^.

## Acknowledgments

We would like to thank Thomas Akam for helpful comments on the manuscript.

## Author contributions

Conceptualization: BM, PD, SWK. Methodology: JLB, BM, WMNM, TEJB, PD, SWK. Investigation: BM, SWK. Visualization: JLB, BM. Funding acquisition: TEJB, PD, SWK. Project administration: PD, SWK. Supervision: PD, SWK. Writing – original draft: JLB, BM. Writing – review & editing: JLB, BM, PD, SWK

## Data and materials availability

All data and code to reproduce figures are available at https://github.com/jamesbutler01/TwoStepExperiment

## Supplementary Figures

**Figure S1.**
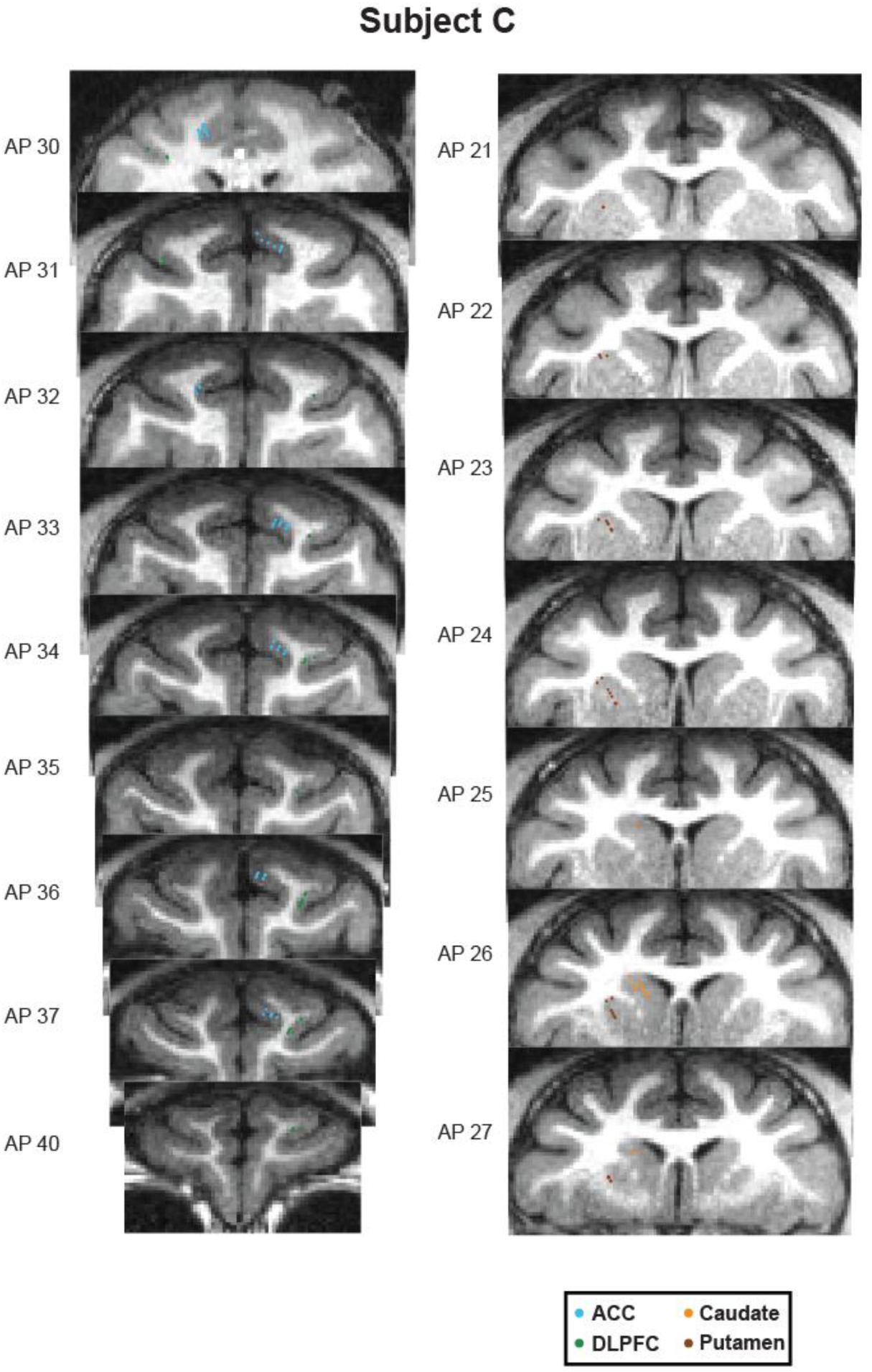
Locations of each neuron recorded from subject C.

**Figure S2.**
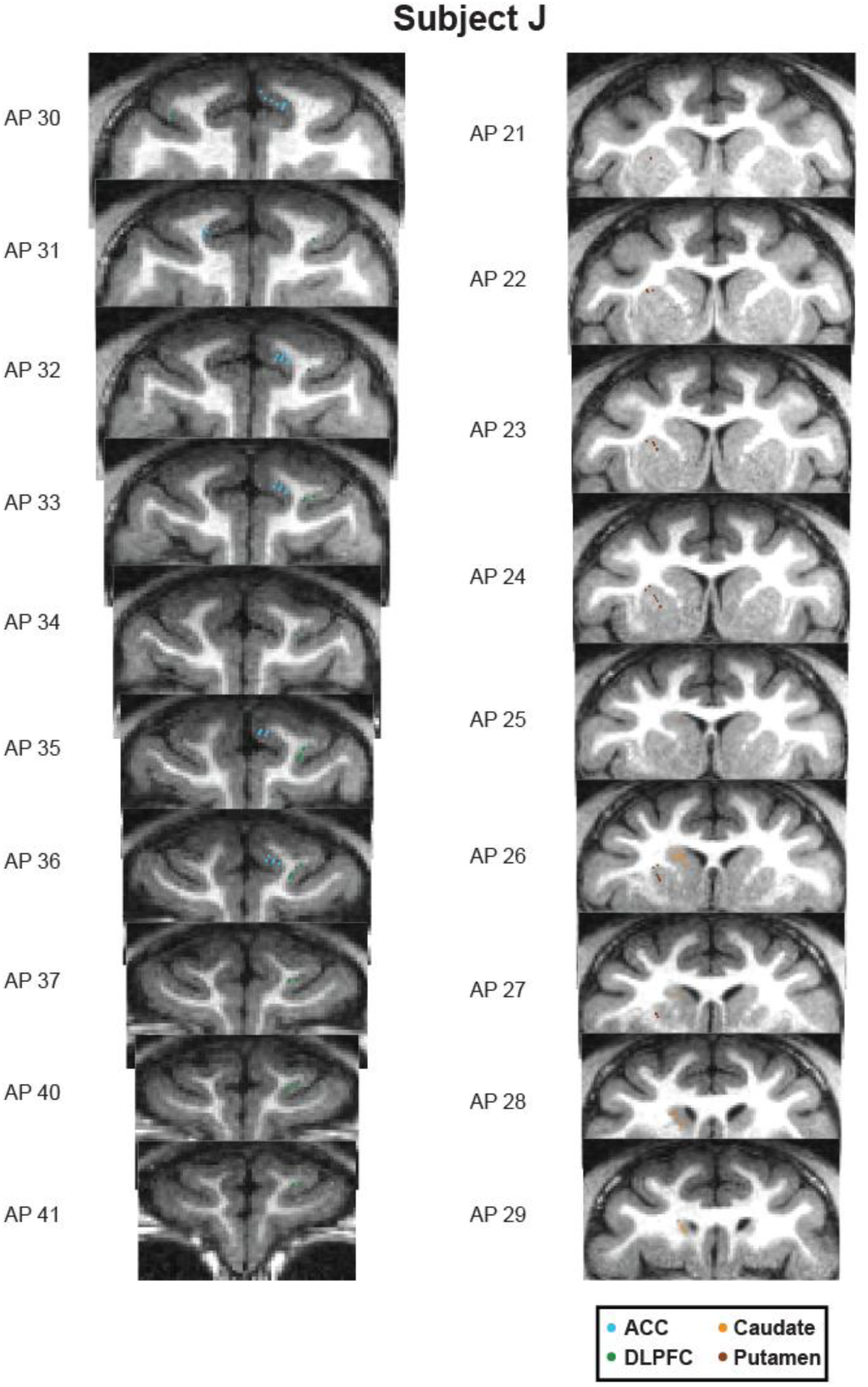
Locations of each neuron recorded from subject J.

**Figure S3.**
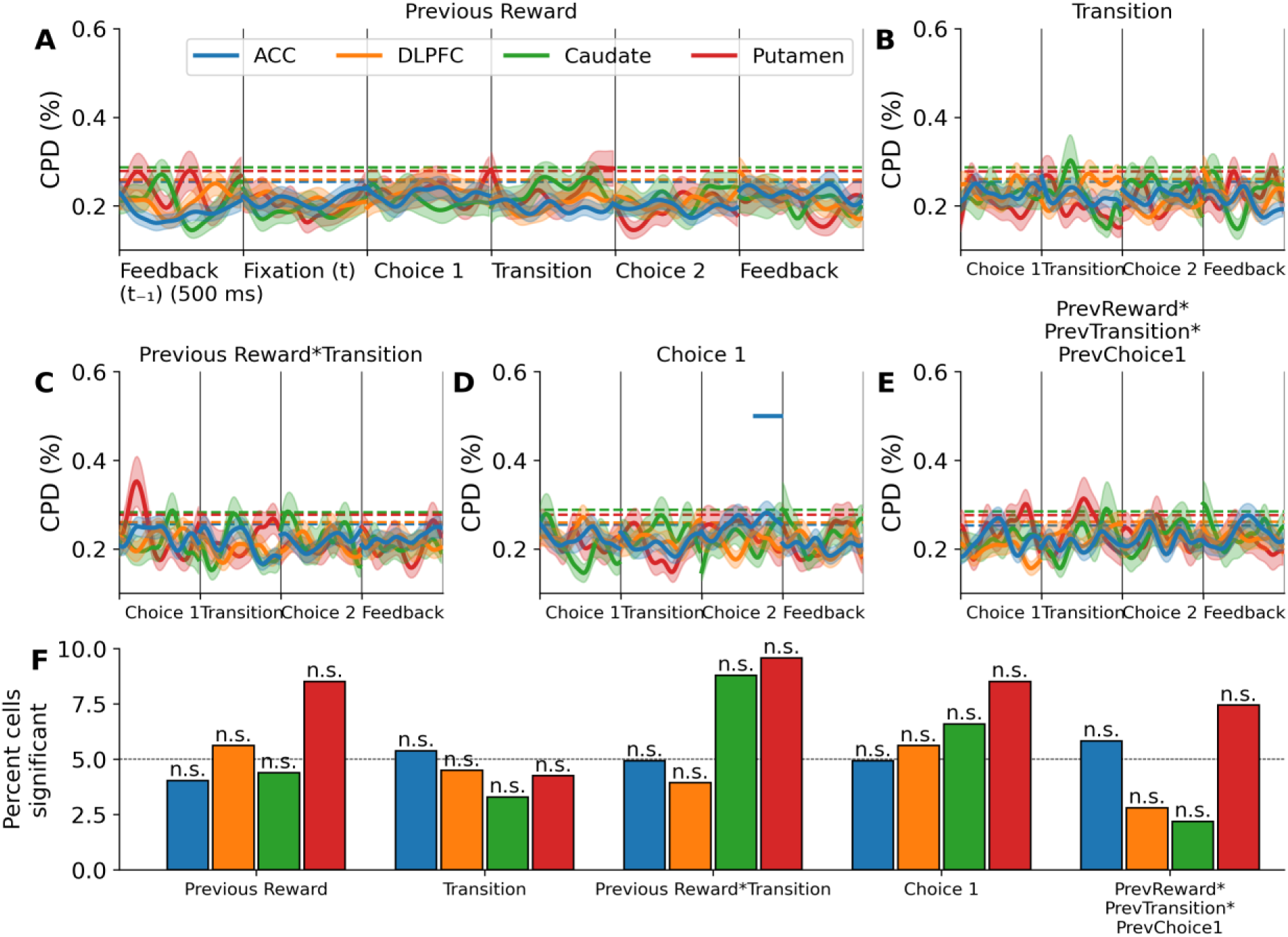
Temporal autocorrelations of neuronal activity did not confound analyses. **A**, To check for the effect of autocorrelations over time we repeated the CPD analysis in Figures 3, 4, and 7 but using trial data from a different session to the neural data. Average coefficient of partial determination (CPD) across neurons in each region for the encoding of the reward received on the previous trial (of a different session). Feedback is shown both for the previous trial (left) and the current trial (right). Solid horizontal lines represent p<0.05 assessed using permutation testing against the null distribution (indicated by dashed horizontal line). Error bars depict standard error of the mean. **B-E**, Same as in **A**, but for the encoding of transition, the interaction of reward and transition, choice 1, and the interaction of previous reward, previous transition and previous choice 1. **F**, The percentage of neurons that were found to significantly encode each parameter in its respective epoch assessed using cluster-based permutation testing (p<0.05). n.s., not significant, binomial test against 0.05 Bonferroni corrected for the 4 regions tested.

**Figure S4.**
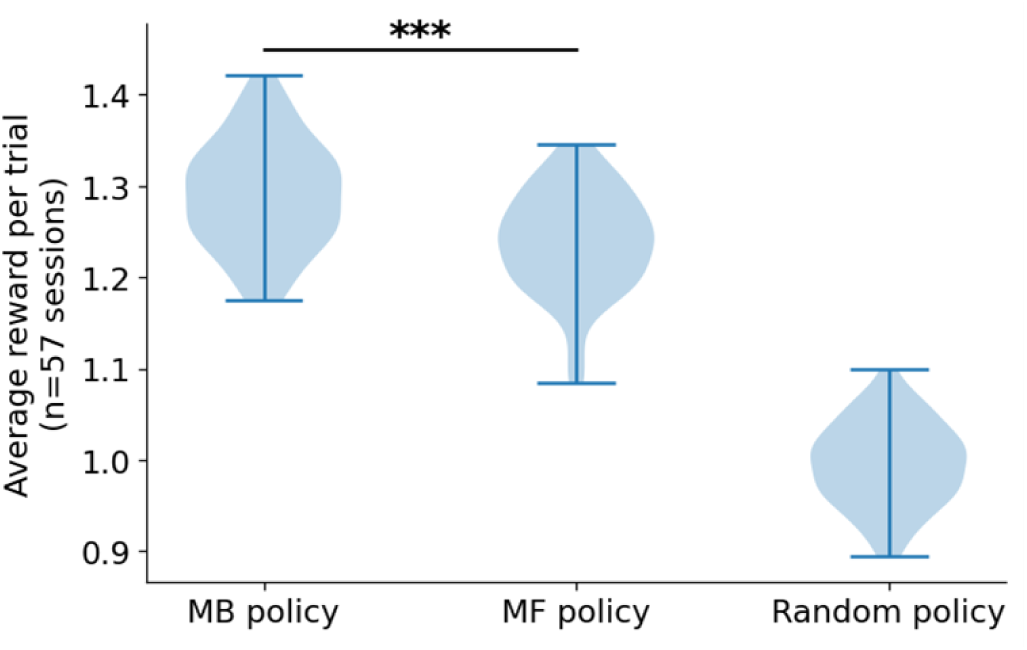
A model-based strategy was best to solve the task. The amount of reward attained by an agent that either always chose the option with the highest MB-derived estimate of value (‘MB policy’) or the highest MF-derived estimate of value (‘MF policy’). This was compared to an agent that chose randomly (‘Random policy’). The average reward per trial over all recording sessions was used (n=57). ***, p<0.001, paired t-test.

**Figure S5.**
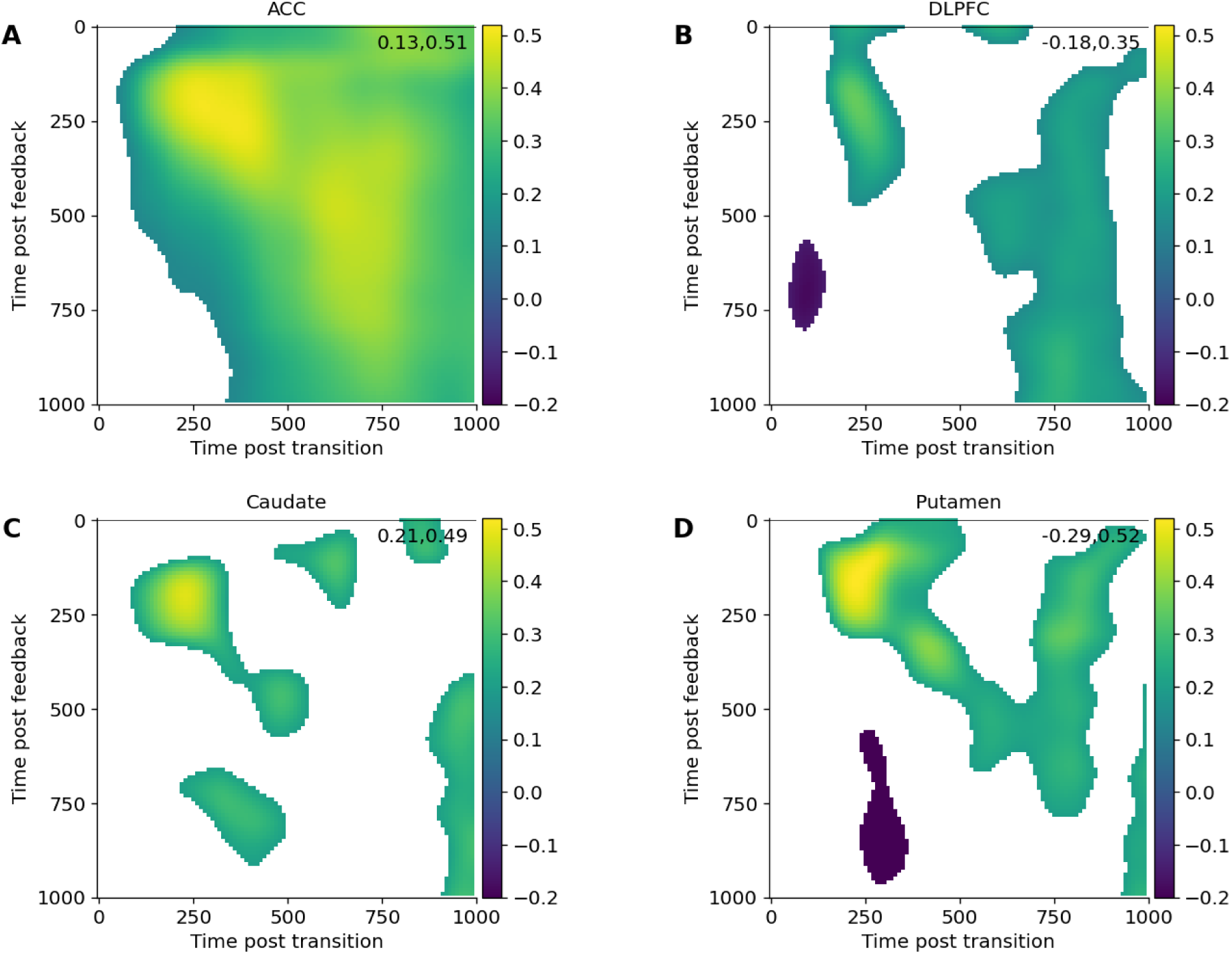
Reward coding and transition coding is correlated. A-D, Correlation of coefficients for transition at the transition epoch (x-axis) with reward at the feedback epoch (y-axis) for ACC, DLPFC, Caudate, and Putamen neurons, respectively. Non-significant points have been removed for clarity (white). Top right numbers indicate the extreme values of each plot (Pearson r).

**Figure S6.**
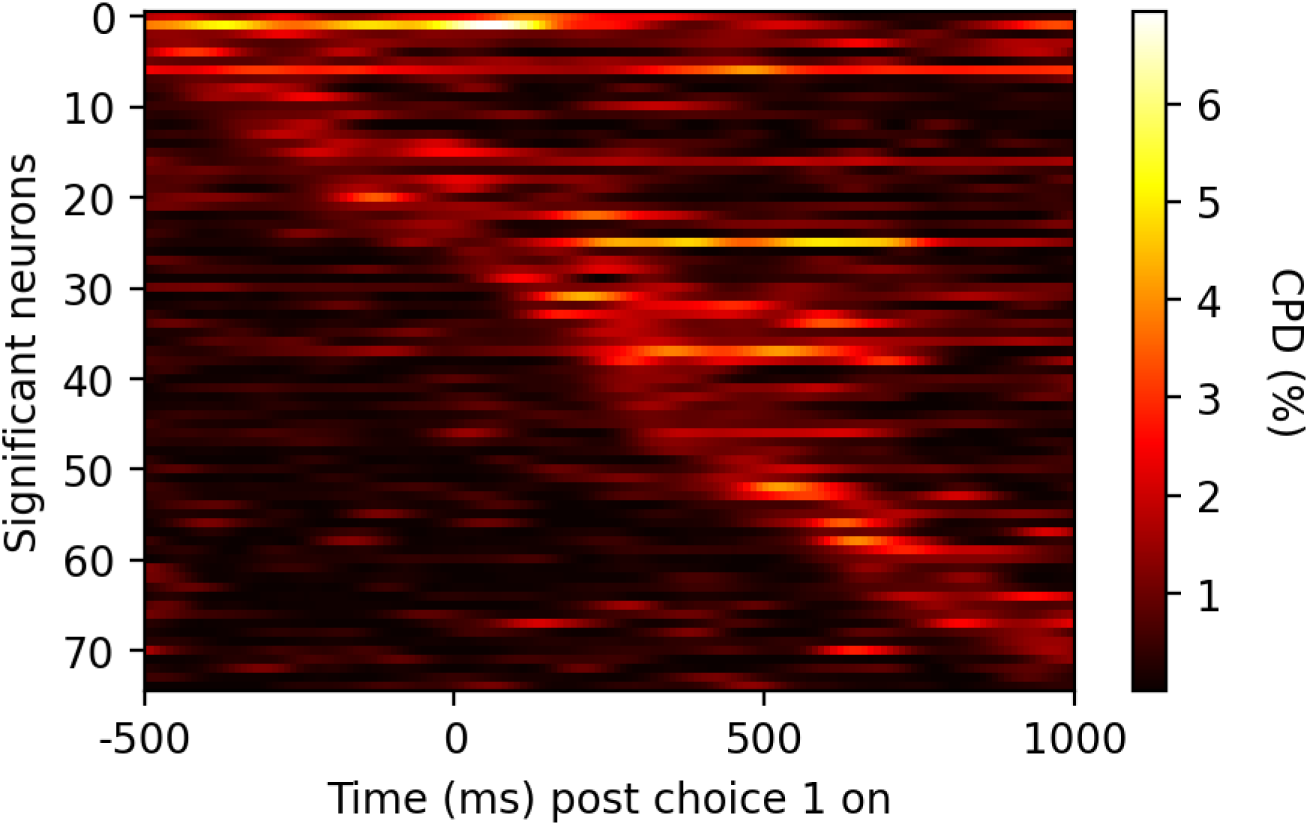
Dynamic coding of model-based Q-values in ACC. Coefficient of partial determination (CPD, z-axis) across significant neurons in ACC for encoding the model-based (MB) derived estimates of the value of each of the choice 1 options (p<0.05, permutation test). Neurons have been sorted by the onset of their coding, revealing a dynamic pattern tiling the entire epoch where different neurons are active at different parts.

**Figure S7.**
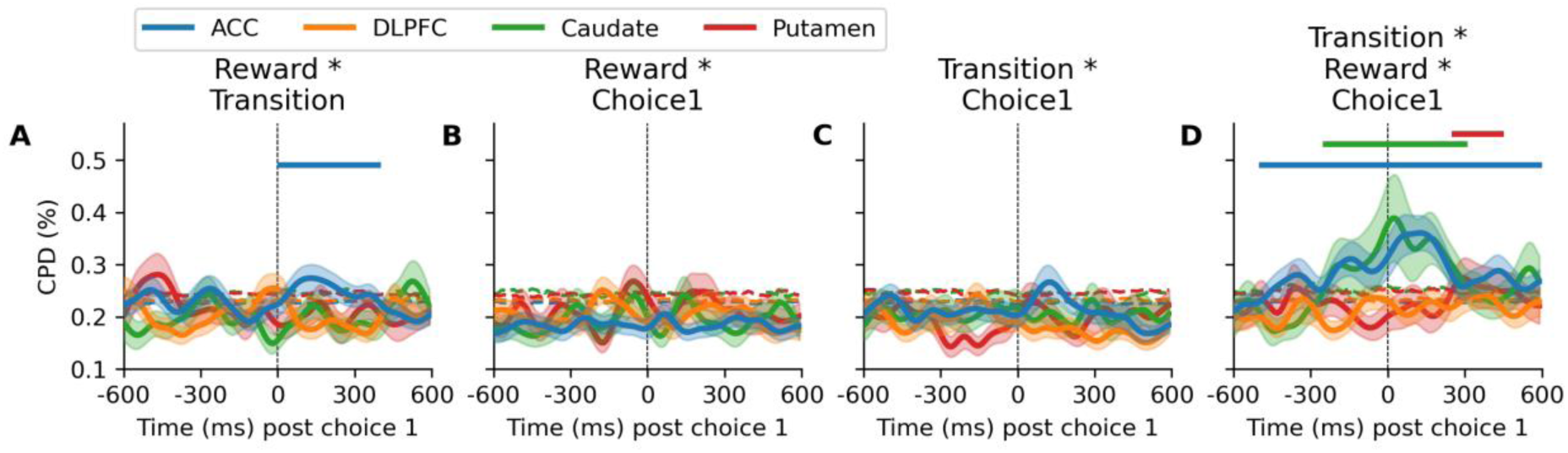
Coding of the interaction of choice 1, transition and reward from the previous trial. **A**, Average coefficient of partial determination (CPD) across neurons in each region for the encoding of the interaction of reward and transition from the previous trial at the time the subjects made their first choice on the next trial. Dashed horizontal lines indicate the 95^th^ percentile of the null distribution and solid horizontal lines indicate periods of significant coding (p<0.05, cluster-based permutation test). Error bars depict standard error of the mean. **B-D**, Same as in A but for the interaction between reward and choice 1, transition and choice 1, and the triple interaction between all three variables.

**Figure S8.**
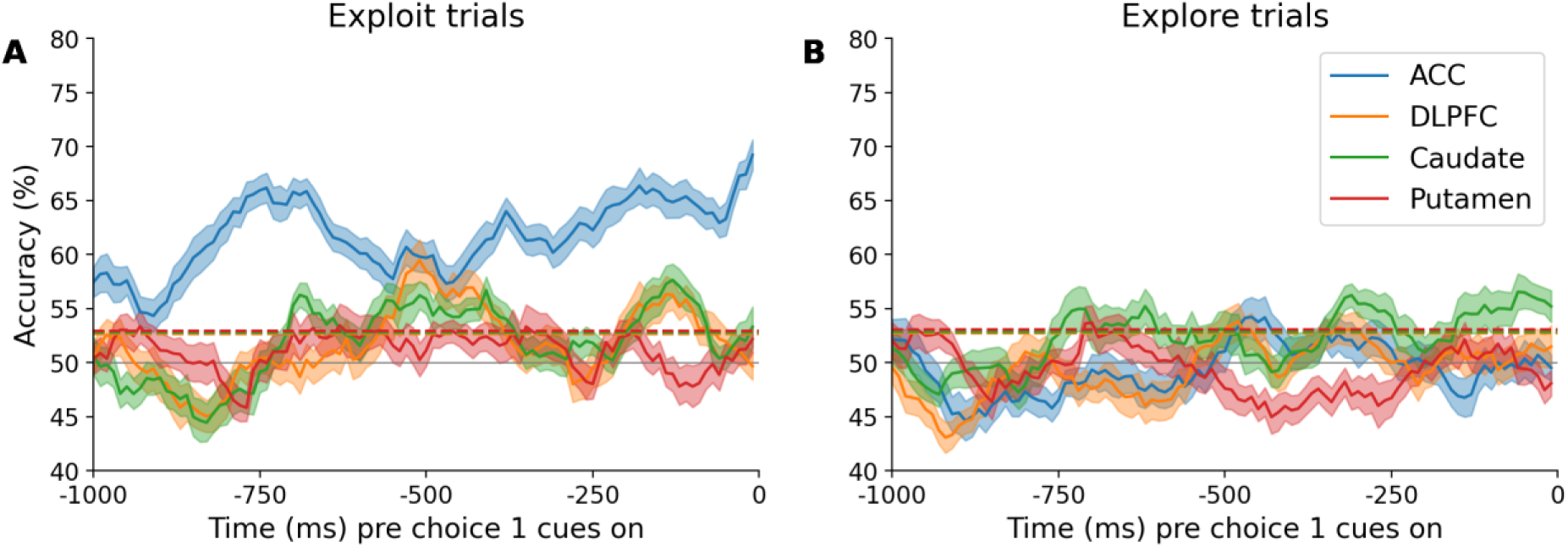
ACC encoded the upcoming choice during the preceding fixation period. **A**, A support vector machine was used to decode from each neural population which cue the monkeys would choose at choice 1 on each trial. Decoding was restricted to trials where they had received a high reward on the previous two trials. Dashed lines represents the 95% confidence interval (permutation test). **B**, Same as in **A** but only for trials following two consecutive low/medium reward trials.

